# Mitochondrial Electron Transport Chain Disruption and Oxidative Stress in Lipopolysaccharide-Induced Cardiac Dysfunction in rats and mice

**DOI:** 10.1101/2025.03.10.642441

**Authors:** Agda Aline Pereira de Sousa, Leonardo da Silva Chaves, Heberty Tarso Facundo

## Abstract

Sepsis, characterized by severe systemic inflammation and an excessive immune response to infection, is frequently triggered by bacterial endotoxins like lipopolysaccharide (LPS) from Gram-negative bacteria. Moreover, sepsis-induced cardiac dysfunction remains a leading cause of mortality. This study aims to elucidate the effects of LPS-induced cardiac injury on mitochondrial damage, oxidative stress, and subsequent cardiac dysfunction. LPS injections (in rats and mice) for three days (1.5 mg/kg) impacted the body weight and increased cardiac TNF-α. Additionally, it decreased mitochondrial complexes I and II activities while complexes III and IV remained unaffected. Disturbed in mitochondrial electron transport chain leads to an increase in reactive oxygen species (ROS). Indeed, LPS treatment significantly increased mitochondrial hydrogen peroxide production, reduced the activity of antioxidant enzymes catalase, superoxide dismutase, glutathione peroxidase, and glutathione reductase activity. This was accompanied by decreased mitochondrial and cytosolic sulfhydryl proteins and parallel increased cellular lipid peroxidation in the presence or absence of Fe^2+^. LPS-treated samples had increased glutathione s-transferase activity, which may be an attempt of the cell to remove toxic lipid peroxidation products. In a more acute Langendorff-perfused rat hearts, LPS infusion (0.5 μg/mL) induced a significant elevation in left ventricular end-diastolic pressure and a decrease in left ventricular developed pressure. These findings elucidate the harmful mitochondrial and oxidative effects of LPS in cardiac tissue and could help the development of targeted therapies to mitigate the adverse effects of sepsis-induced cardiac dysfunction.

## Introduction

Sepsis is a pathological condition initiated by a dysregulated immune response to a widespread infection. In its most severe form, known as septic shock, the systemic inflammatory response leads to multiple organ dysfunction and a significant increase in mortality rates. This exacerbated inflammatory and immune response is characterized by the excessive release of pro-inflammatory mediators, frequently triggered by bacterial endotoxins, such as lipopolysaccharides (LPS) [1,2]. Sepsis-induced myocardial dysfunction is a reversible myocardial dysfunction associated with sepsis that is characterized by contractile dysfunction, dilation of ventricular chambers, and a reversible decline in left ventricular ejection fraction [3,4]. Despite extensive research, the complex pathological mechanisms of sepsis, involving mitochondrial dysfunction, oxidative stress, dysregulated calcium handling, and immune system dysregulation, pose significant challenges in developing effective therapeutic approaches that address all these mechanisms simultaneously.

Cardiomyocytes require a constant supply of ATP, as the heart is highly dependent on oxidative phosphorylation to maintain its contractile function [5,6]. During sepsis, the electron transport chain becomes defective [7–12] and produces an excessive amount of reactive oxygen species (ROS) [9,10,13] . Therapies aimed at protecting mitochondria from high ROS generation ameliorates the cardiac tissue damage [14]. ROS are highly reactive chemical molecules formed from O_2_ including superoxide anion (O_2_^•-^), hydrogen peroxide (H_2_O_2_), and hydroxyl radical (•OH). Studies have shown that cardiomyocytes treated with LPS exhibit a reduction in respiratory activity in their mitochondria, followed by increased mitochondrial oxidative stress [9,10]. Furthermore, LPS can induce the expression of NADPH oxidases (NOXs), which produce O_2_ ^•−^, through the transfer of electrons from NADPH [15]. The first line of defense against excessive ROS is antioxidant enzymes (superoxide dismutase – both cytosolic and mitochondrial) that dismutate O_2_ ^•−^ to H O (a less reactive ROS), glutathione peroxidase and catalase break down H_2_O_2_, glutathione reductase (responsible for reducing GSSG back to GSH), and glutathione S-transferase, which neutralizes final oxidative products such as lipid peroxidation products with glutathione.

Cardiac dysfunction caused by sepsis is a significant factor contributing to mortality and morbidity rates, particularly among patients admitted to intensive care units [3]. Key aspects of this dysfunction include impaired ventricular ejection fraction and compromised contractile function of the heart, primarily characterized by a reduced pumping capacity of the left ventricle (LV) and, in some cases, right ventricle (RV) failure due to pressure overload from LV failure [16]. This left ventricular depression is a well-established manifestation in patients with severe sepsis [1,4]. Mitochondria constitute approximately one-third of the cellular volume, supplying the high ATP levels necessary to sustain cardiomyocyte contractility and relaxation [17]. Sepsis disrupts ATP levels in cardiomyocytes [18]. Thus, a comprehensive understanding of the basic molecular and pathological events underlying sepsis is crucial for advancing therapeutic strategies. Importantly, there is still uncertainty in the literature regarding specific mitochondrial functional and oxidative changes occurring in septic cardiomyopathy. Points to be clarified or detailed include: the effects on mitochondrial complexes, the parallel production of ROS, the activities of the main antioxidant enzymes and the consequent damage to macromolecules (lipids and proteins). Here, we provide a side-by-side study of all these parameters in mouse and rat models. We report that LPS-induced sepsis for three days depresses mitochondrial complex I and II activity. This is accompanied by high H_2_O_2_ production by cardiac isolated mitochondria fed with substrates from complex I and II. We also report that LPS negatively affects antioxidant enzymes (catalase, SOD, Glutathione peroxidase and reductase) activity. As a consequence, this leads to an elevated level of oxidative damage to lipids and proteins. Finally, besides more sustainable changes we also found that LPS induces diastolic dysfunction in the rat heart.

## Materials and methods

### Animals and Ethics

We used male Wistar rats and Swiss mice. All procedures were approved by the Institutional Animal Experimentation Ethics Committee of the Universidade Federal do Cariri (Protocol Number 0002/2023). All experiments were performed according to the Guide for the Care and Use of Laboratory Animals published by the National Institutes of Health. Animals were divided into 2 groups: 1. LPS group was injected with LPS 1.5 mg/kg from Escherichia coli O111:B4 (Sigma Aldrich Code L-2630) diluted in saline and 2. Control group was injected with saline for 3 days.

### Mitochondrial Isolation

Male rats were anesthetized with a mixture of ketamin (100 mg/kg of body weight) and xilazin (10 mg/kg of body weight). Then, their hearts were immediately removed, washed, and finely chopped into small pieces in an ice-cold buffer consisting of 300 mM sucrose, 10 mM K^+^ HEPES buffer, and 1 mM K^+^ EGTA at pH 7.2. The tissue samples were incubated with protease type I (Sigma-Aldrich) for 10 minutes, followed by two washes with ice-cold buffer containing 1 mg/mL BSA to inhibit protease activity. Subsequently, the tissue was manually homogenized using a glass homogenizer (potter). An aliquote (∼0.5 ml) of total homogenate was used to assay for thiobarbituric acid reactive substances (TBARS) as described later in this paper. Nuclei and intact cellular debris were removed by centrifugation at 1200 g for 5 minutes. The resulting supernatant was centrifuged again at 9400 g for 10 minutes to obtain a mitochondrial pellet, which was then resuspended in a minimal volume of buffer (approximately 100-150 μL). The supernatant (cytosolic fraction) was used for biochemical analysis such as antioxidant enzymes. Mitochondria were maintained on ice throughout the procedure.

### Mitochondrial H_2_O_2_ quantification

To assess H_2_O_2_ levels under each experimental condition, we incubated isolated mitochondria from mice and rats (n = 4) with substrates (2 mM malate plus 4 mM glutamate which feeds complex I or 2 mM succinate to feed electrons to complex II), amplex red (50 μM), and horseradish peroxidase (1 U/mL) for 30 minutes at 37 °C in the dark. The reaction buffer consisted of 100 mM KCl, 10 mM HEPES, 2 mM MgCl_2_, and 2 mM KH_2_PO_4_ at pH 7.2 (adjusted with KOH). After centrifugation at 9400 x g for 2 minutes, the supernatant was measured at 560 nm using a spectrophotometer zeroed with a blank reaction containing amplex red and horseradish peroxidase but no mitochondrial sample. Mitochondrial H_2_O_2_ production was quantified by using a standard curve generated with known concentrations of H_2_O_2_ and expressed as μmol/mg protein.

### Mitochondrial complexes activity

Mitochondrial complexes activity was detected following the protocol described in [19]. First, mitochondria (200 μg) were lized in hypotonic buffer composed of 25 mM de KH_2_PO_4_ and 2.5 mM MgCl_2_ pH 7.2 by submitting mitochondrial suspension in two cycles (5 minutes each) of freeze thaw in – 80°C. We measured complex I spectrophotometrically at 340 nm in 50 mM K Phosphate pH 7.4 in the presence of 50 µM decylubiquinone. The reaction was initiated by NADH 100 μM. We followed the absorbance decay due to the consumption of NADH by complex I. Then, we inhibited complex I using 4μM rotenone and follow absorbance of the inhibited complex I. Complex I specific activity is calculated as nanomoles/min/mg protein using the molar extinction coefficient for NADH of 6.2 mM^−1^ cm^−1^. The final activity is the decay in the absence rotenone minus the decay in the presence of rotenone (NADH decay rotenone insensitivity).

We tested the activities of complexes II, III, and IV by observing changes in the absorbance of 10 μM reduced cytochrome C at 550 nm [19]. Briefly, oxidized cytochrome C was reduced using sodium dithionite in phosphate buffer (pH 7.0), prepared on the day of the experiment. The reduction of cytochrome C was confirmed by calculating the ratio of absorbance values at 550 nm to 565 nm, with a ratio greater than 6 considered appropriate for use. First, we detected complex IV activity by adding lysed mitochondria (200 μg) and monitoring the decay in absorbance of reduced cytochrome C. Complex IV was inhibited using 0.3 mM KCN. At this stage, 8 μM rotenone was also added to inhibit complex I. Next, we tested the combined activities of complexes II and III by adding 5 mM succinate, which donates electrons to complex II, then to ubiquinone, and finally to complex III, resulting in the reduction of cytochrome C. To stop this reaction, we added 5 mM malonate to inhibit complex II. To test the activity of the antimycin A-sensitive complex III, we added 50 μM decylubiquinol. This reduced ubiquinone transfers electrons to complex III, which in turn reduces cytochrome C.

### Catalase activity

We measured catalase activiy by following the changes in H2O2 absorbance at 240 nm. Supernatants of homogenized samples (as described earlier) were added to the reaction media containing 50 mM H_2_O_2_ in 100 mM of phosphate buffer (pH 7.4). The reaction was calibrated to start at absorbances close to 0.5. Changes in absorbance at 240 nm were recorded for 2 minutes. Catalase activity was calculated as milliunits of catalase per milligram protein.

### Superoxide dismutase (SOD) activity

SOD activity was assayed in a reaction medium containing 0.1 mM EDTA, 13 mM L-methionine, and 75 mM nitro blue tetrazolium (NBT) in potassium phosphate buffer (pH 7.8). The 2 μM riboflavin was added and exposed uniformly to an unfiltered white light for 5 minutes. The developed blue color due to NBT reduction was measured at 560 nm. SOD activity was expressed as U/mg of protein. One unit is the amount of enzyme required to inhibit the reduction of NBT by 50%.

### Glutathione peroxidase activity assay

Glutathione peroxidase activity was determined by the decrease in NAD(P)H absorbance at 340 nm for 5 minutes. We used H_2_O_2_ as a substrate. First the cellular extract was added to a buffer containing 50 mM potassium phosphate, 0.5 mM EDTA, 0.2 mM NADPH (Sigma Aldrich), 1 mM GSH (Sigma Aldrich) and 0.2 U/mL Glutathione reductase purified from S. cerevisiae (Sigma Aldrich), pH 7.0. Hydrogen peroxide was added to initiate the reaction after an initial period of incubation of 5 minutes. Glutathione peroxidase activity is expressed as Units/mg protein.

### Glutathione S-transferase

We measured glutathione S-transferase activity spectrophotometrically at 340 nm. The reaction mixture contains an aliquot of the cardiac homogenate (cytosolic fraction) in potassium phosphate buffer pH 6.5, 1 mM GSH, and 1 mM 1-chloro-2,4-dinitrobenzene (CDNB), which is used as a substrate. The enzyme conjugates GSH with CDNB forming 2,4-dinitrophenil S-glutathione (DNP-SG) which was monitored for 10 minutes at room temperature at 340nm. Enzymatic activity was calculated by GST by using the extinction molar coefficient of DNP-SG = 9.6 mM^−1^.cm^−1^ and expressed in mU/mg protein.

### Glutathione reductase

Glutathione reductase activity was determined by following the reduction of GSSG to 2GSH by the enzyme which also transforms NAD(P)H to NAD^+^. Glutathione reductase activity is proportional to the decay of NAD(P)H, which was monitored at 340 nm. Glutathione reductase was calculated by using the extinction molar coefficient of NAD(P)H (6.2 mmol/L^−1^) cm^−1^ and expressed in mU/mg protein.

### Protein thiol content

We measured protein sulfhydrils (thiols) content using 0.2 mM DTNB (prepared in PBS plus 1 mM EDTA) incubated in the dark with homogenate supernatants (prepared as described above) for 30 minutes at room temperature. DTNB reacts with free sulfhydrils generating a yellow product (TNB) which absorbs at 412 nm. The protein thiol content was calculated based on the molar extinction coefficient of TNB (14500 M^−1^cm^−1^) and reported as moles of TNB per milligram protein. The blank had DTNB with no protein and was submitted to the same incubation.

### Thiobarbituric acid reactive substances (TBARS)

TBARS were estimated in cardiac homogenates spectrophotometrically at 540 nm. Samples were tested in the absence (basal peroxidation levels) and 30 µM Fe^2+^ (for 25 minutes) to further stimulate lipid oxidation and test for homogenates susceptibility to *in vitro* oxidation. Then, 1.2 ml TBARS reagent (15% (w/v) TCA, 0.38% (w/v) TBA), and heated at 95 °C for 15 min. After cooling, the tubes were centrifuged for 5 min at 2,000 x g and the supernatants read at 540 nm. Results were expressed as mol TBARS/mg protein.

### Enzyme-linked immunosorbent assay (ELISA) for TNF-α

Cardiac homogenates were used for TNF-α detection using ELISA. Levels of TNF-α in heart homogenates (control or LPS-treated groups) were determined by mouse-specific ELISA kit (sigma-aldrich code RAB0477) according to the manual. Data are expressed relative to control.

### Heart perfusion experiments

Male Wistar rats were used for experiments LPS perfusion in isolated hearts. Briefly, rats were anesthetized and sacrificed, the hearts removed and aorta cannulated (less than 3 minutes for the cannulation). The cannulated hearts were perfused (at a constant flow of 12 mL min^−1^) with oxygenated (gas mixture bubbled with 95% O_2_, 5% CO_2_) Krebs-Henseleit buffer containing (in mM) 118 NaCl, 25 NaHCO_3_, 1.2 KH_2_PO_4_, 4.7 KCl, 1.2 MgSO_4_, 1.25 CaCl_2_, 10 glucose, and 10 HEPES, pH 7.4, at 37°C in a Langendorff apparatus (AD Instruments). To measure hemodynamic function, we inserted a latex balloon (filled with water) connected to a pressure transducer (Powerlab/8SP, AD Instruments). After stabilization of the heart was injected 0.5 μg/mL LPS in oxygenated krebs buffer. Control hearts were perfused with the same volume of PBS.

### Cardiac hemodynamic parameters

We recorded the pressures of the heart in a computer-based data acquisition system (PowerLab 8S/P with LabChart 4 software, AD Instruments). Using the program Lab Chart reader (version 8.1) to calculate the left ventricular systolic pressure (LVSP), left ventricular end-diastolic pressure (LVEDP), and left ventricular developed pressure (LVDP, LVDP = LVSP-LVEDP).

### Statistical analysis

Statistical analysis was conducted using Graphpad Prism software. Data are presented as mean ± S.E.M. Student’s t test was used to test for statistical differences. P < 0.05 was considered statistically significant.

## Results

### LPS induces loss of weight and upregulation of TNF-α

Animals injected with LPS for 3 days showed a significant weight loss (Fig. 1A) 2 days after the start of the injections. This was also accompanied by an increased upregulation of TNF-α in cardiac homogenates (Fig. 1B). These results suggest that our protocol or LPS injections effectively induce sepsis.

**Fig. 1.**
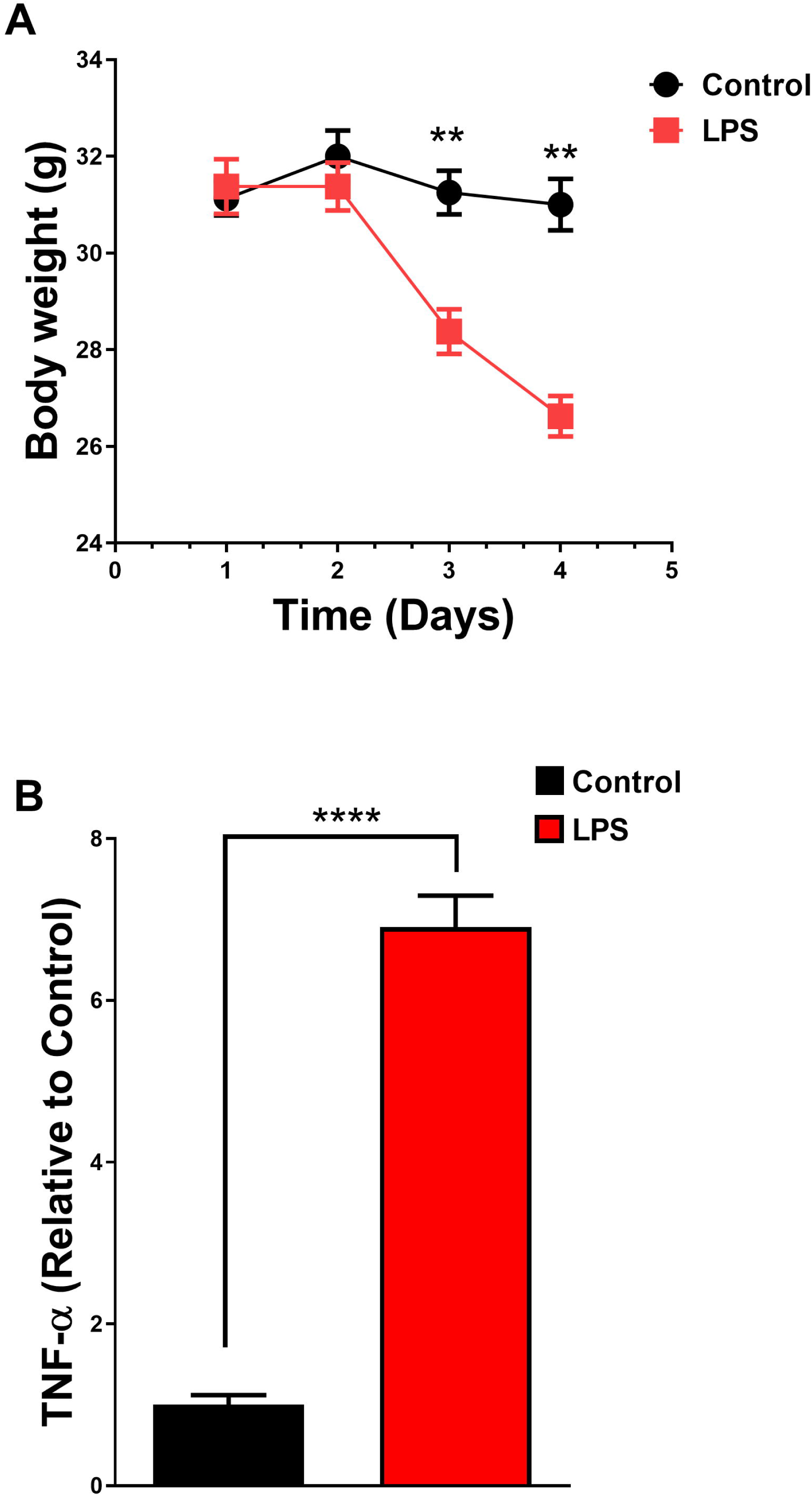
LPS-induced sepsis is accompanied by loss of weight and cardiac inflammatory signaling. A, body weight of mice during LPS treatment. B, TNF-α levels in cardiac homogenates of LPS-treated mice. n = 4-5. **, P < 0.01, ****, P < 0.0001.

### Mitochondrial respiratory chain complexes I and II are downregulated by LPS-induced sepsis in vivo

Mitochondria are the powerhouse of the cell participating in important processes such as calcium homeostasis, produce about 95% of the ATP used by the heart, are important in several signaling pathways and may trigger cell death [17]. Knowing the importance of mitochondria in several processes we sought to directly measure the activity of all four complexes of the electron transport chain in LPS-induced sepsis *in vivo*. We determined the activity of complex II+III, III, and IV using the oxidation/reduction state of added reduced cytochrome C (redCytC). RedCytC absorbs at 550 nm. As shown in Fig. 2A (scheme), and representative traces (control - Fig. 2B and sepsis Fig. 2C) the addition of redCytC to mitochondrial lysates leads to its oxidation since complex IV removes the electron from redCytC and transfers it to oxygen present in the reaction medium (Fig. 2A, IV). LPS-induced sepsis did not change complex IV activity in mice (Fig. 2D) or rat (Fig. supplementary 1A) cardiac mitochondrial lysates. From that point of the curve, we inhibited complex IV using KCN and complex I with rotenone. Then, we fed the mitochondrial electron transport chain with succinate Fig. 2A (scheme II+III), Fig. 2B (control) and C (sepsis -representative traces). In control samples succinate induced an elevation of redCytC (complex II+III activity) in a manner sensitive to malonate (Fig. 2B). Remarkably, LPS-treated samples were insensitive to succinate (Fig. 2C). This suggests a defect in mitochondrial electron transport between complex II or III or both. Indeed, averages ± standard errors show a decreased activity of complex II+III in LPS-treated cardiac samples (Fig. 2D). To pinpoint the defect, we fed complex III directly with decylbuquinol (a reduced form of decylubiquinone which feeds electrons to complex III, Fig. 2A). Using this protocol, we found that control and LPS-septic mitochondrial lysates had preserved activities of antimycin-sensitive complex III (Fig. 2B, Fig. 2C for representative traces, and D for averages ± SEM). These results were also reproduced in rat tissue (Supplementary Fig. 1A). These results suggest that LPS-induced sepsis leads to a reduction in mitochondrial complex II electron transport capacity. Finally, we tested the activity of mitochondrial complex I by testing the rotenone-sensitive NADH decay in control and LPS-treated samples (Fig. 3A, 3B, 3C). As shown in Fig. 3D (mice) and Supplementary Fig. 1B (rat), sepsis *in vivo* led to a strong reduction in mitochondrial complex I activity. Interestingly, the NADH consumption in LPS-treated samples is rotenone insensitive (Fig. 3C). These results strongly suggest that LPS-induced sepsis disturb mitochondrial electron transport chain.

**Fig. 2.**
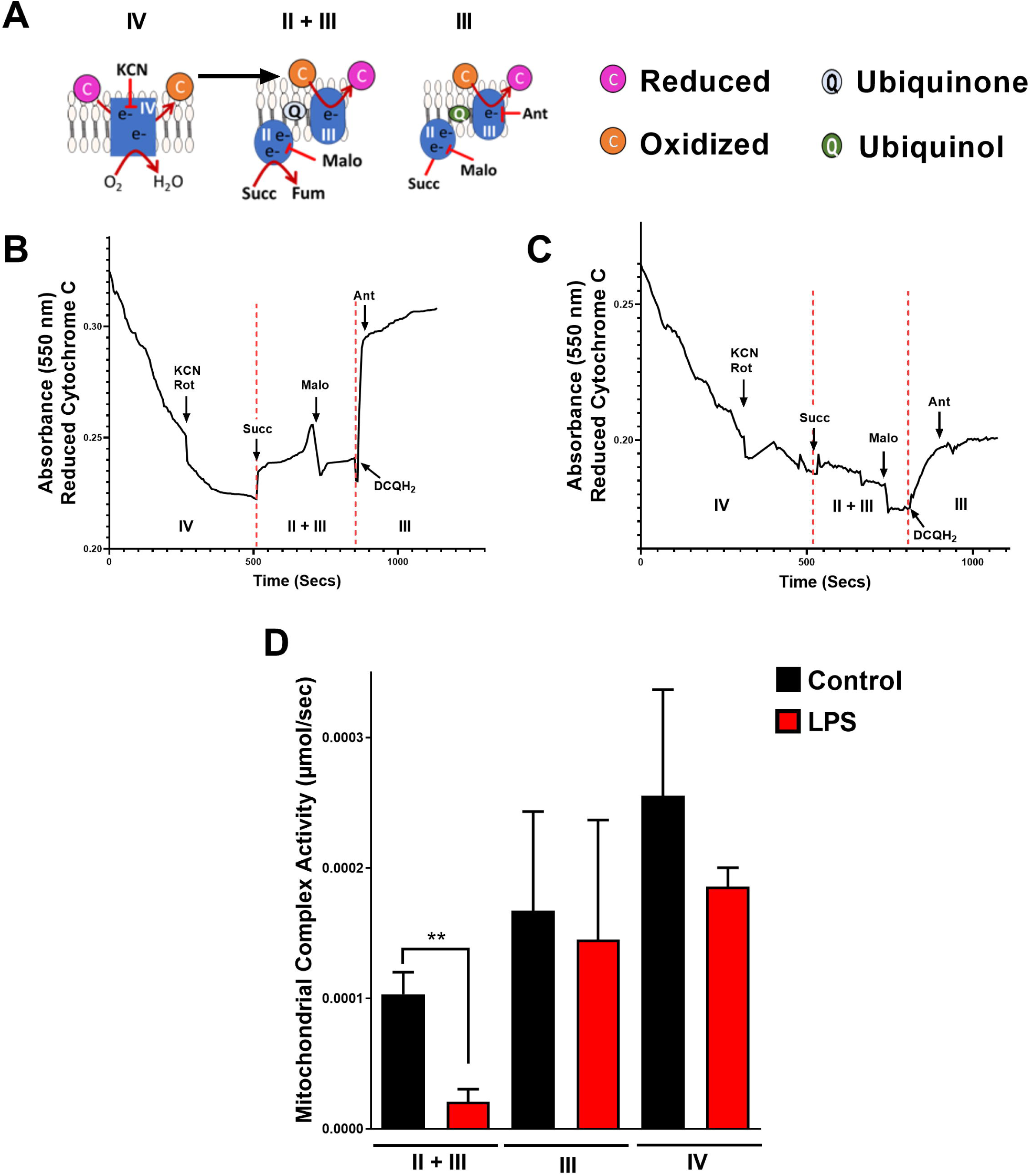
Impaired cardiac mitochondrial electron complex II activity in LPS-treated animals. A, scheme of electron transport activity protocol. Representative traces of electron transport test for complex II+III, III and IV from controls (B) and LPS-treated mice samples (C). Panel D shows average ± SEM of electron transport activity (complex II+III, III and IV) of at least four repetitions using different mice cardiac mitochondria preparations. * P < 0.05.

**Fig. 3.**
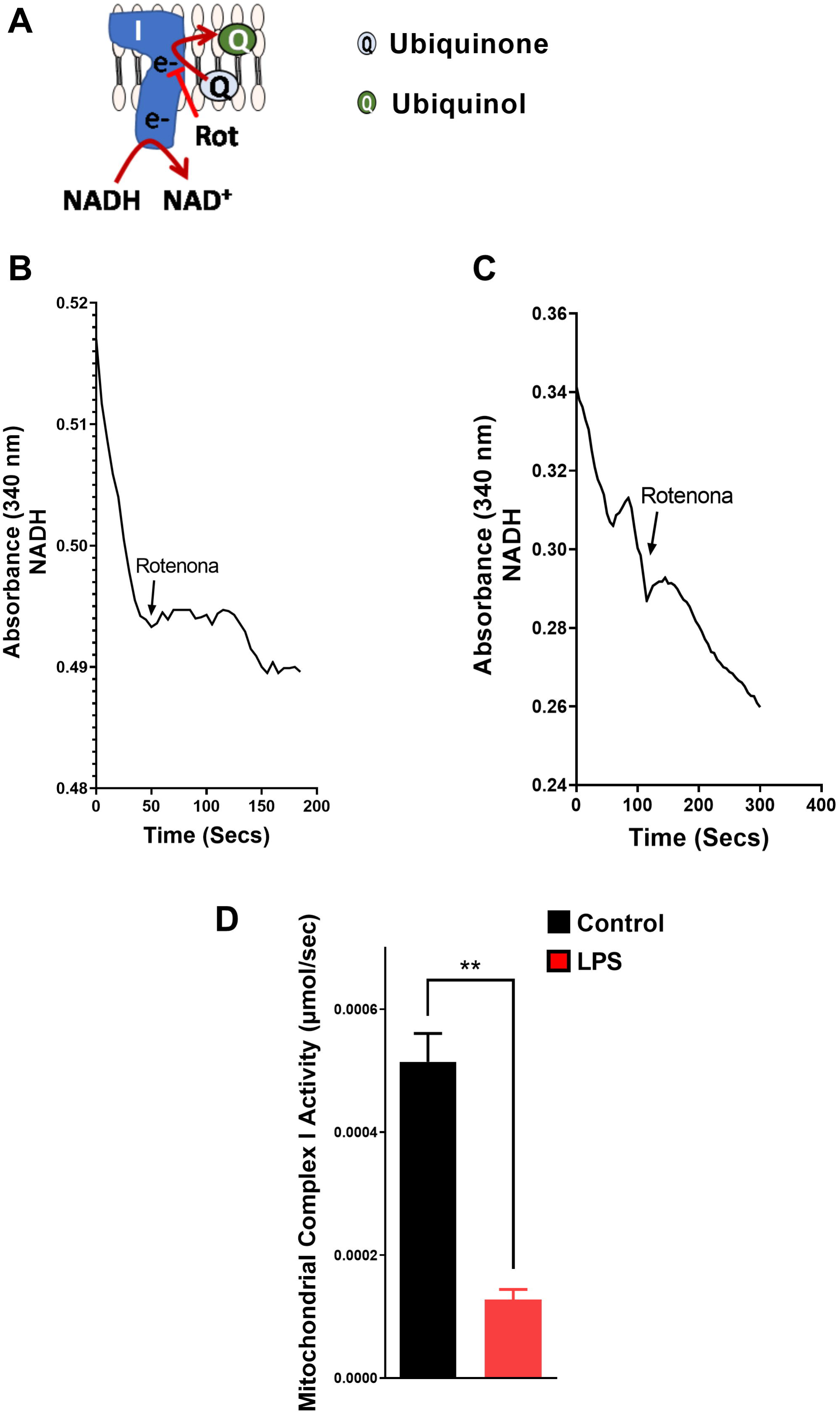
LPS-induced sepsis affects cardiac mitochondrial electron complex I activity. A, scheme of electron transport activity protocol. Representative traces of rotenone-sensitive electron transport test for complex I from controls (B) and LPS-treated samples (C). Panel D shows average ± SEM of electron transport activity (complex I) of at least four repetitions using different cardiac mitochondria preparations. * P < 0.05.

### LPS-induced sepsis in vivo leads to ROS production, repression of antioxidant enzyme activity, and oxidative damage

The majority of studies consistently report a reduction in complex I activity or impaired complex I-driven mitochondrial respiration during sepsis [7–9,11,12], which results in the overproduction of reactive oxygen species (ROS) [9,10,13]. Corroborating these findings, our data also demonstrate decreased complex I activity (Fig. 3), accompanied by elevated H_2_O_2_ production in mitochondria utilizing complex I substrates (malate/glutamate, Supplementary Fig. 2). Additionally, although less consistently reported, the dysregulation of complex II is a significant concern [8–10,12]. Notably, complex II plays a pivotal role in mitochondrial metabolism, and its deficiency is linked to cardiac pathogenesis [20]. To understand the consequences of complex II dysfunction in cardiac mitochondria triggered by LPS-induced sepsis, we probed for ROS formation (H_2_O_2_) in mitochondria fed with complex II (succinate). This setup tests for any reverse electron transport triggering ROS formation or complex II, III derangements [21,22]. As shown in Fig. 4, mitochondria from LPS-treated mice (Fig. 4A) or rats (Fig. 4B) produced higher levels of H_2_O_2_ when stimulated with complex II substrate succinate, thus supporting that the derangements in mitochondrial electron transport (seen in Fig. 2, and 3) lead to oxidative stress in LPS-treated animals. Consistent with this, LPS-treated mice (Fig. 5) and rats (Supplementary Fig. 3) also had lower activity of antioxidant enzymes catalase (Fig. 5A), superoxide dismutase (Fig. 5B), glutathione peroxidase (Fig. 5C) and glutathione reductase (Fig. 5D). Conversely, LPS-treated samples had higher activity of glutathione S-transferase (Fig. 5E). This enzyme is highly active against lipid peroxidation products and may be upregulated in the presence of highly concentrated peroxidative products [23]. This suggests that lipid peroxidation may be high in LPS samples. Indeed, increased lipid peroxidation (TBARS levels) was observed in LPS-treated samples with or without stimulation with Fe^2+^ (Fig. 6A and supplementary 4). Finally, we also found that LPS-treated samples had lower protein sulfhydryl (thiol – another marker of oxidative imbalance) in mitochondria and cytosol (Fig. 6B and C). Together, these results suggest that sepsis induces mitochondrial oxidative imbalance contributing to lipid and protein oxidation.

**Fig. 4.**
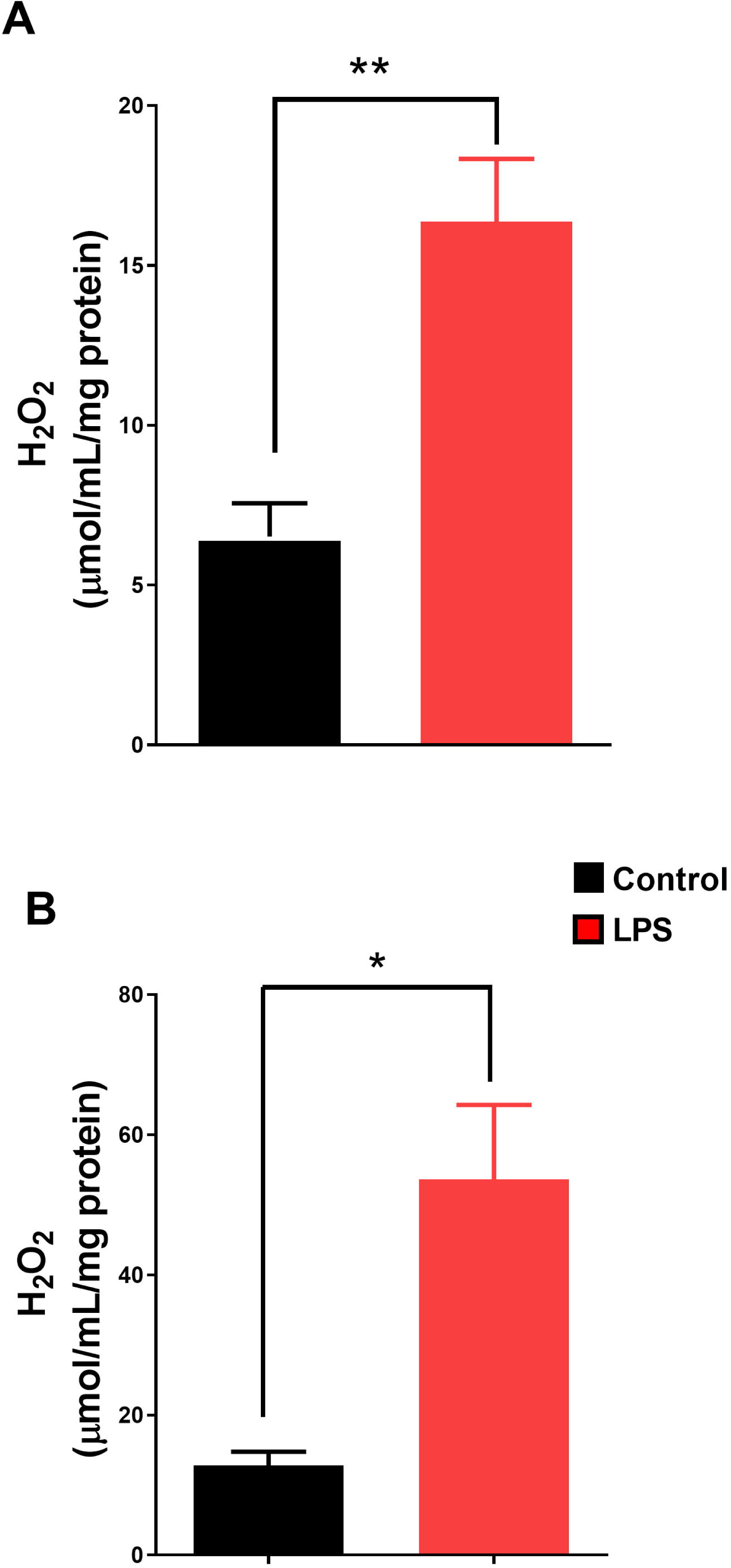
LPS-induced sepsis increases mitochondrial H_2_O_2_ production. Isolated mice (A) or rats (B) heart mitochondria (0.2 mg/mL, n = 4) were incubated in 100 mM KCl, 10 mM K-HEPES (pH 7.2), 2 mM MgCl_2_, 2 mM KH_2_PO_4_, and 1 μg/mL oligomycin, in the presence of 2 mM succinate. * P < 0.05.

**Fig. 5.**
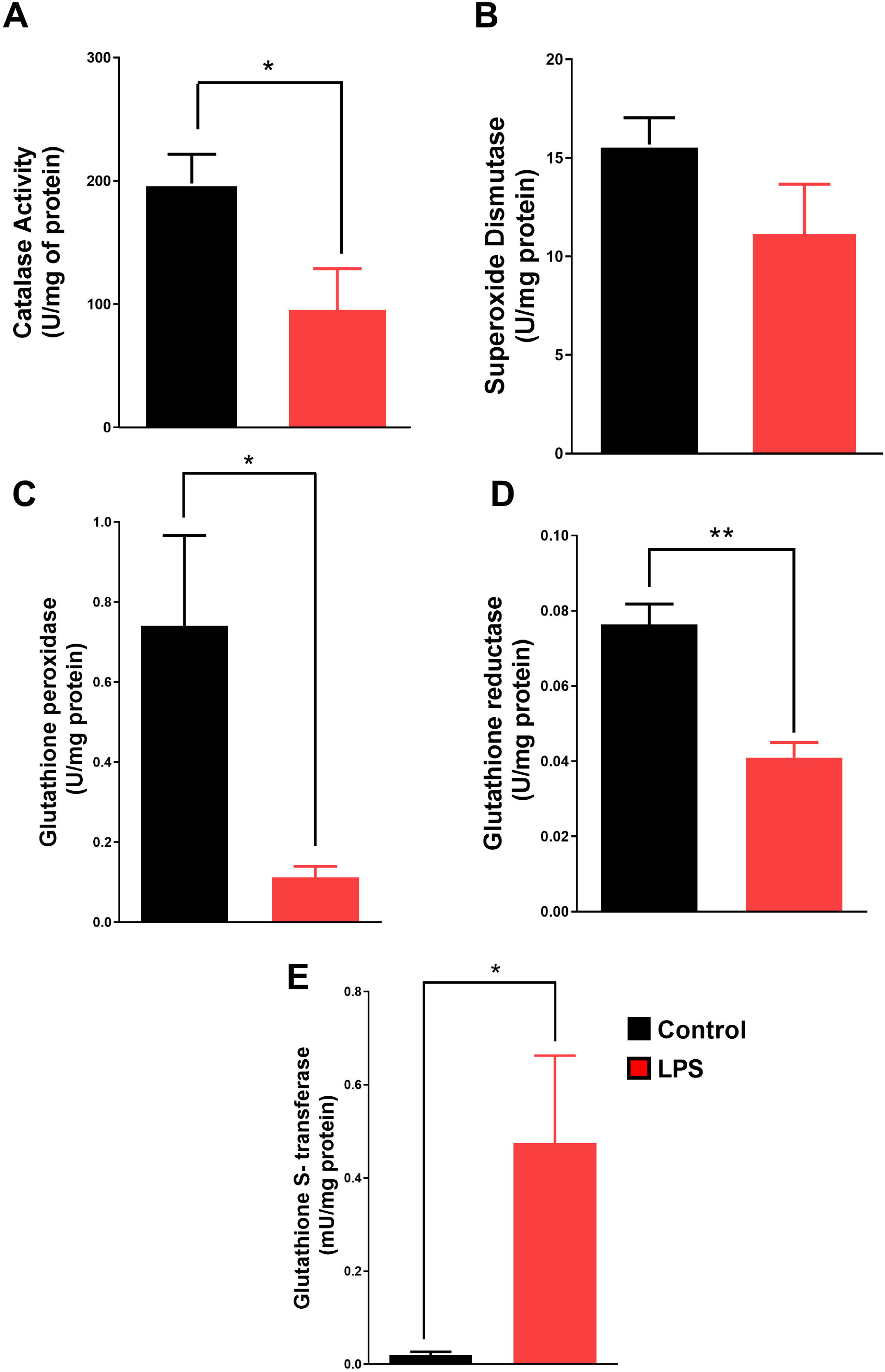
Sepsis depresses cardiac antioxidant enzymes activity while upregulating glutathione S-transferase activity. A, catalase activity. B, superoxide dismutase activity. C, glutathione peroxidase activity. D, glutathione reductase activity. E, glutathione S-transferase activity. (n = 4-5). * P < 0.05, ** P < 0.01.

**Fig. 6.**
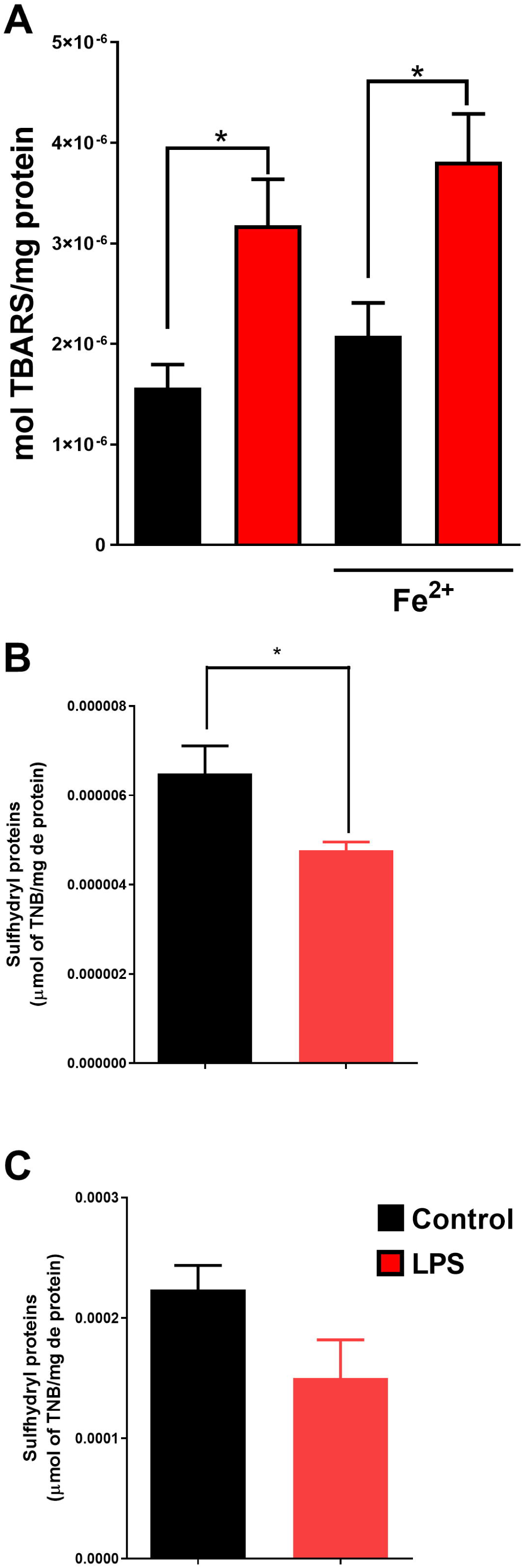
Sepsis induces lipid and protein oxidation in mice cardiac tissue. A, Lipid peroxidation (TBARS) increases in LPS-induced sepsis in the presence or absence of Fe^2+^. Mitochondrial (B) and cytosolic (C) protein sulfhydryl levels. (n = 4-5). * P < 0.05, ** P < 0.01.

### LPS-induced cardiac hemodynamic effects

The cardiac tissue needs mitochondrial fitness to contract and transport ions. The heart consumes large quantities of energy and possesses low energy storage. Mitochondria produce more than 95% of the ATP in the myocardium, besides participating in metabolism, calcium homeostasis, and redox signaling [17,24]. It is germane to hypothesize that LPS could acutely change cardiac functioning prior to any sustained mitochondrial and redox signaling impairment as seen in the results above. To test the direct effect of LPS on cardiac function, we asked whether LPS could affect the cardiac contraction/relaxation cycle. Indeed, LPS treatment (0.5 μg/mL) on *ex vivo* Langendorff heart preparation for 30 mins impaired cardiac hemodynamic parameters. The most affected functional parameter was LVEDP. After direct contact (30 minutes) with LPS the cardiac tissue becomes hypercontracted which is seen by the increase in LVEDP (Fig. 7A). Conversely, LVSP was not affected (Fig. 7B). The elevation in LVEDP seen in LPS treated hearts negatively impacted the LVDP (Fig. 7C). These results suggest that acutely injected LPS induces a diastolic dysfunction in the rat heart.

**Fig. 7.**
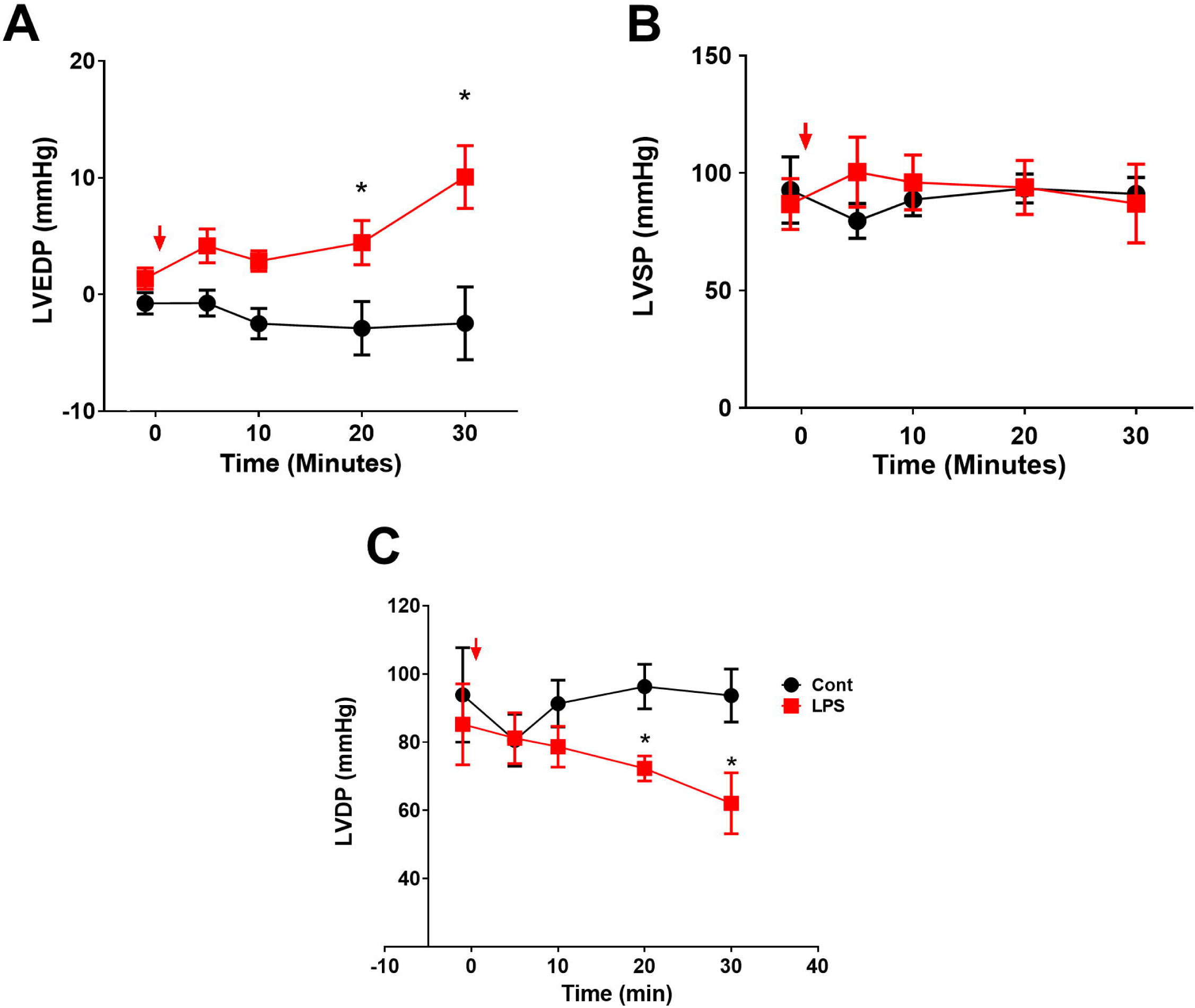
Sepsis induces rat cardiac contractile dysfunction. A, Left ventricular end diastolic pressure (LVEDP). B, Left ventricular systolic pressure. C, Left ventricular developed pressure (LVDP) calculated as LVSP-LVEDP. Arrows indicate LPS injection. (n = 4-5). * P < 0.05.

## Discussion

Septic cardiomyopathy induces organ failure and increases the risk of death in patients. Sustained sepsis-induced myocardial damage is characterized by myocardial contractile depression [3,4]. It is well accepted that mitochondrial function may be compromised in septic cardiomyopathy [7–11], but the literature is not clear about the specific changes and how mitochondria are damaged during sepsis. Given that mitochondrial damage is central to myocardial injury during sepsis, it is crucial to understand the fundamental mechanisms underlying sepsis. This study demonstrates that in rat and mouse models of LPS-induced sepsis, multiple pathological effects are observed ultimately resulting in impaired cardiac performance. LPS-induced sepsis chronically manifests as disrupted mitochondrial electron transport (with decreased activity of complexes I and II). This disruption is associated with an imbalance in cellular enzymatic antioxidants, increased ROS generation, and elevated lipid/protein oxidation.

To sustain high contractility and relaxation cardiomyocytes need high levels of ATP with mitochondria as the main source. Indeed, cardiomyocytes are densely packed with mitochondria that occupy about one-third of the volume of cardiac myocytes [17]. In Fig. 2 and 3 we found that mitochondria from LPS-treated animals had impaired cardiac mitochondrial complex I and II activity. LPS treatment has been shown to reduce state 3 (ADP stimulated) mitochondrial oxygen consumption in complex I fed substrates. Differences in complex II fed mitochondria are not consistent [8,10,12], or the rate of O_2_ consumption increased in the presence of LPS [9]. Transient reductions in complex I activity and respiration are also reported with a transient decrease in complex II fed respiration [10]. In agreement with our finding some have demonstrated a decrease in activity of complex I and II rats [11]). There is uncertainty whether complex II is decreased in sepsis since this change is not so consistent, as said. Mitochondrial respiratory complex II (succinate dehydrogenase), is very important to mitochondrial metabolism since it links the tricarboxylic acid cycle (TCA cycle, or Krebs cycle) with the electron transport chain [25]. Mitochondrial complex II deficiency leads to cardiac pathogenesis [20]. Therefore, deficiency in complex II activity might be responsible for sustained cardiac metabolic derangements and poor cardiac performance. We recognize that inhibition of complex II is an important tool to block reverse electron transport and avoid cardiac damage induced by ischemia/reperfusion [26]. The point here is that LPS (contrary to other reports [8–10]) triggers a more sustainable loss of complex II activity what may impact metabolism and mitochondria impacting cell function. Others also found decreased complex IV activity in LPS-induced sepsis which was accompanied by mitochondrial swelling, and cristae disruption [11]. Our experiments showed no differences in complex IV activity in either model mice or rat cardiac tissue. It is expected that deficiency in mitochondrial electron transport decreases ATP formation and enhances ROS generation which triggers damage. These results raise the possibility that mitochondrial complexes defect could be a direct result of oxidative stress to mitochondrial membranes, protein subunits assembly disruption or poor import into mitochondria. All these changes would impact mitochondrial complex functioning and finally impact oxidative phosphorylation [7,11]. We may exclude assembly disruption or poor import into mitochondria since activity of complex III and IV were not affected by LPS treatment (Fig. 2). We raise the hypothesis that mitochondrial complexes are, firstly, impacted by inflammatory signaling which will lead to leakage of electrons to O_2_ increasing ROS which will impact mitochondrial structure (ex. protein thiol oxidation) and homeostasis and, finally, will lead to poor cardiomyocyte performance.

It is clear that LPS-induced sepsis leads to mitochondrial derangements in electron transport system (Fig. 2 and 3). The immediate functional consequence of mitochondrial complexes failure is electron leakage to the available O_2_ forming ROS. Physiologically, ROS are removed by a set of antioxidants which transform, breakdown and neutralize them. If produced in high levels and not removed properly these ROS may lead to oxidation of cellular components. A key strength of our study is the assessment of the impact of LPS on mitochondrial electron transport, ROS generation, antioxidant enzymes and cellular components (lipids and proteins) oxidation in the same samples making the results correlatable. Importantly, these were verified in mice and rats that are the two most used animal models [27]. Our results point to an increased mitochondrial leakage of electrons forming ROS (H_2_O_2_, Fig. 4). These ROS are not removed properly since LPS-induced sepsis also presents low levels of antioxidant enzymes (catalase, SOD, glutathione peroxidase and reductase – Fig. 5). This will result in oxidative stress which causes damage to macromolecules and impair cardiac function. Complex I is a major site for mitochondrial ROS formation both forward or reverse electron transport [21]. The adverse effect of complex I reduced activity would be a large electron leakage forming ROS. It is reasonable to hypothesize that mitochondrial-mediated ROS generation leads to ROS damage to macromolecules in the immediate surrounding of complex I, making mitochondria a site for ROS formation and also a target. Lipids such as cardiolipin (which is particularly rich in unsaturated fatty acids) which is important for mitochondrial electron transport including complex I activity [28], could be a ROS target. Here, we did not test for direct cardiolipin oxidation in our model but we show that LPS induced a higher level of overall lipid peroxidation (Fig. 6) with or without stimulation by Fe^2+^. The lipid peroxidation products are toxic and upregulates oxidative stress and cell death. Importantly, we found that LPS-treated samples had an upregulation of glutathione S-transferase activity (Fig. 5). We believe this upregulation is related to an attempt by the cell to counteract the toxic lipid peroxidation-derived aldehyde products (ex. 4-Hydroxy-2-trans-nonenal - 4HNE) through its conjugation to GSH [23].

A major question in the field relates to how cardiac performance is disturbed in sepsis. There is robust evidence for cardiac left ventricular depression in sepsis [1,4,16]. We believe that LPS-triggered inflammatory and Ca^2+^ signaling [29] or oxidative stress triggered by NADPH induces [15] induces a poor cardiac performance that will be accompanied by the sustained changes seen in our paper and by others (mitochondrial electron transport chain defects and oxidative stress). These mitochondria and oxidative stress changes will lead to a potent vicious cycle that worsens organ performance even further, leading to adverse outcomes. A limitation of our study is that we are not able to measure cardiac performance, mitochondrial and oxidative stress changes *in vivo* after each LPS injection. In summary, several approaches, including mitochondrial complexes activity, mitochondrial ROS production and antioxidant enzyme activities were used to assess the effects of LPS in cardiac tissue during sepsis.

In conclusion, this research highlights the harmful impact of LPS-induced cardiac damage on mitochondrial integrity and functioning, resulting in elevated oxidative stress and triggering and maintaining cardiac dysfunction.

## DECLARATIONS

### Ethical aproval

All procedures were approved by the Institutional Animal Experimentation Ethics Committee of the Universidade Federal do Cariri under protocol number (0002/2023).

### Authors’ contributions

Heberty T Facundo: Conceptualization, Investigation, Writing – original draft Writing – review & editing. Agda Aline Pereira de Sousa, Leonardo da Silva Chaves: Investigation, Methodology. Heberty T Facundo wrote the paper, which was critically reviewed by all authors. All authors read and approved the final manuscript.

### Availability of data and materials

Data will be made available on reasonable request.

## Acknowledgements

The authors acknowledge the technical assistance of Anna Lidia Nunes Varela, Iuliana Marjory Martins Ribeiro, and Antônio F. R. Santos.

---

**Supplementary Fig. 1:**
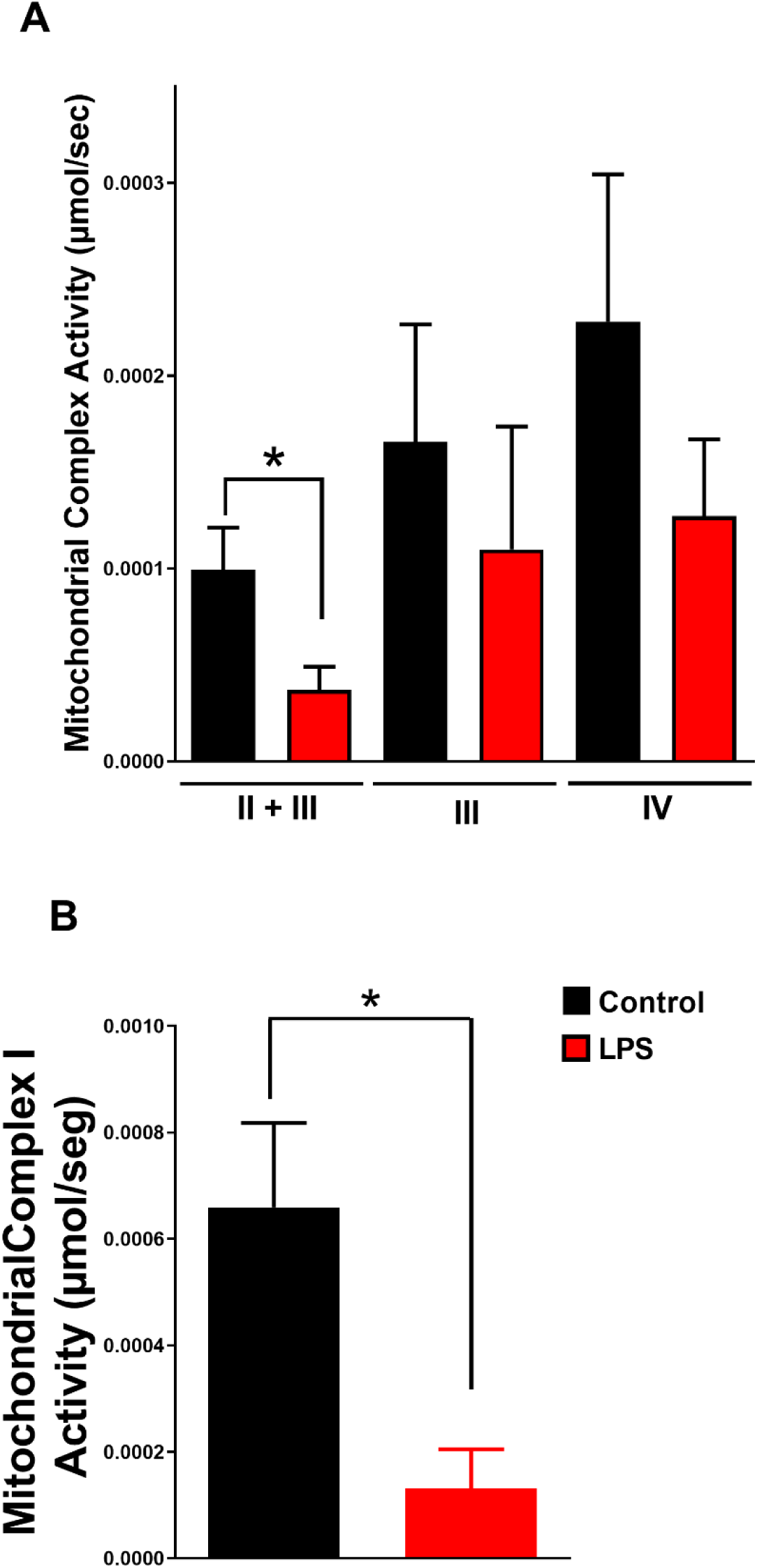
mitochondrial electron transpor chain activity in LPS-treated rats. A, Complex II+III, III and IV from controls and LPS-treated rats samples. B, Complex I activity. Graphics show the average ± standard error of at least four repetitions using different rats cardiac mitochondria preparations. * P < 0.05.

**Supplementary Fig. 2:**
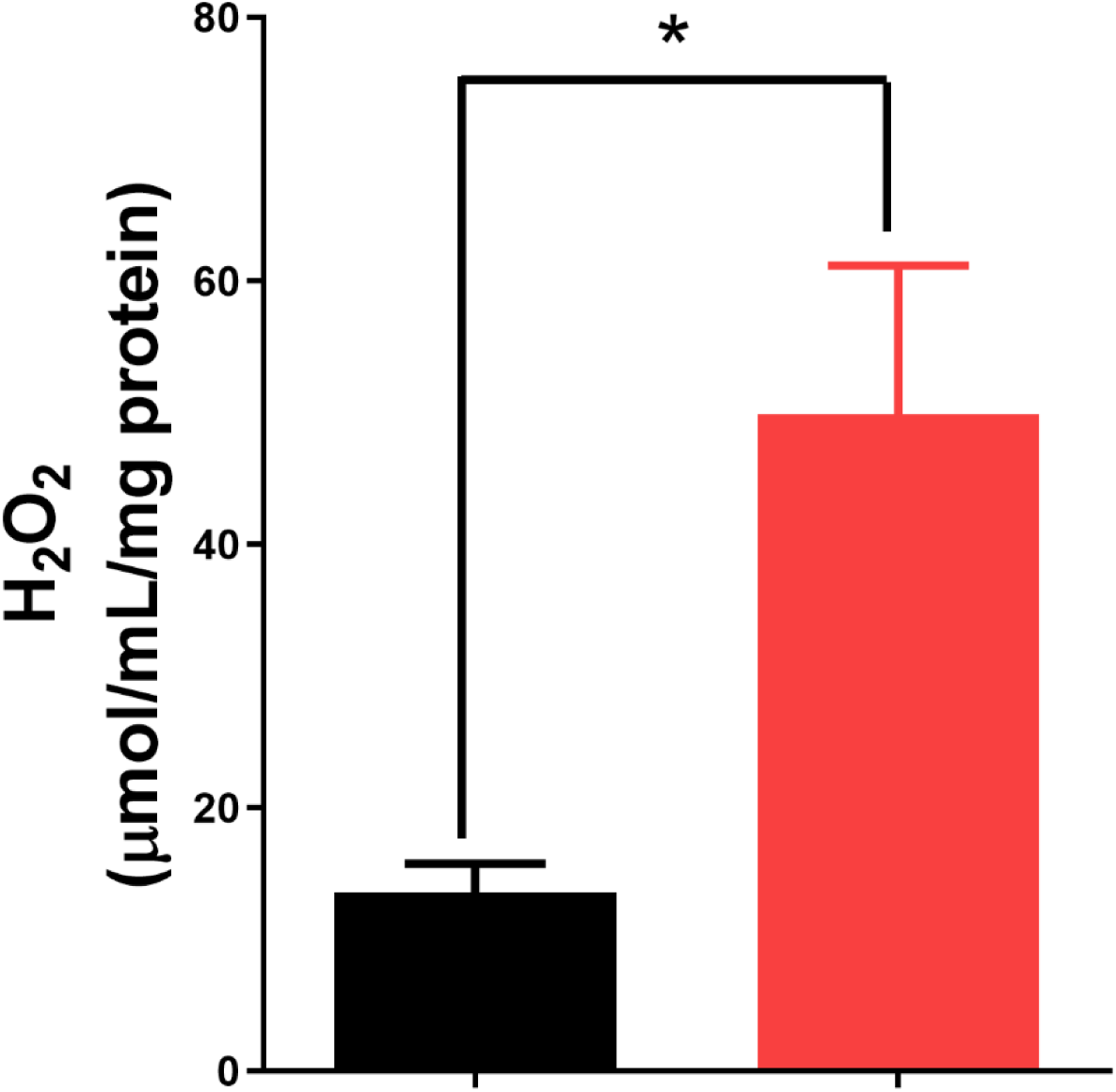
LPS-induced sepsis increases mitochondrial complex I-derived H_2_O_2_ production. Isolated rat heart mitochondria (0.2 mg/mL, n = 4) were incubated in 100 mM KCl, 10 mM K-HEPES (pH 7.2), 2 mM MgCl_2_, 2 mM KH_2_PO_4_, and 1 μg/mL oligomycin, in the presence of 2 mM malate plus 4 mM glutamate. * P < 0.05.

**Supplementary Fig. 3:**
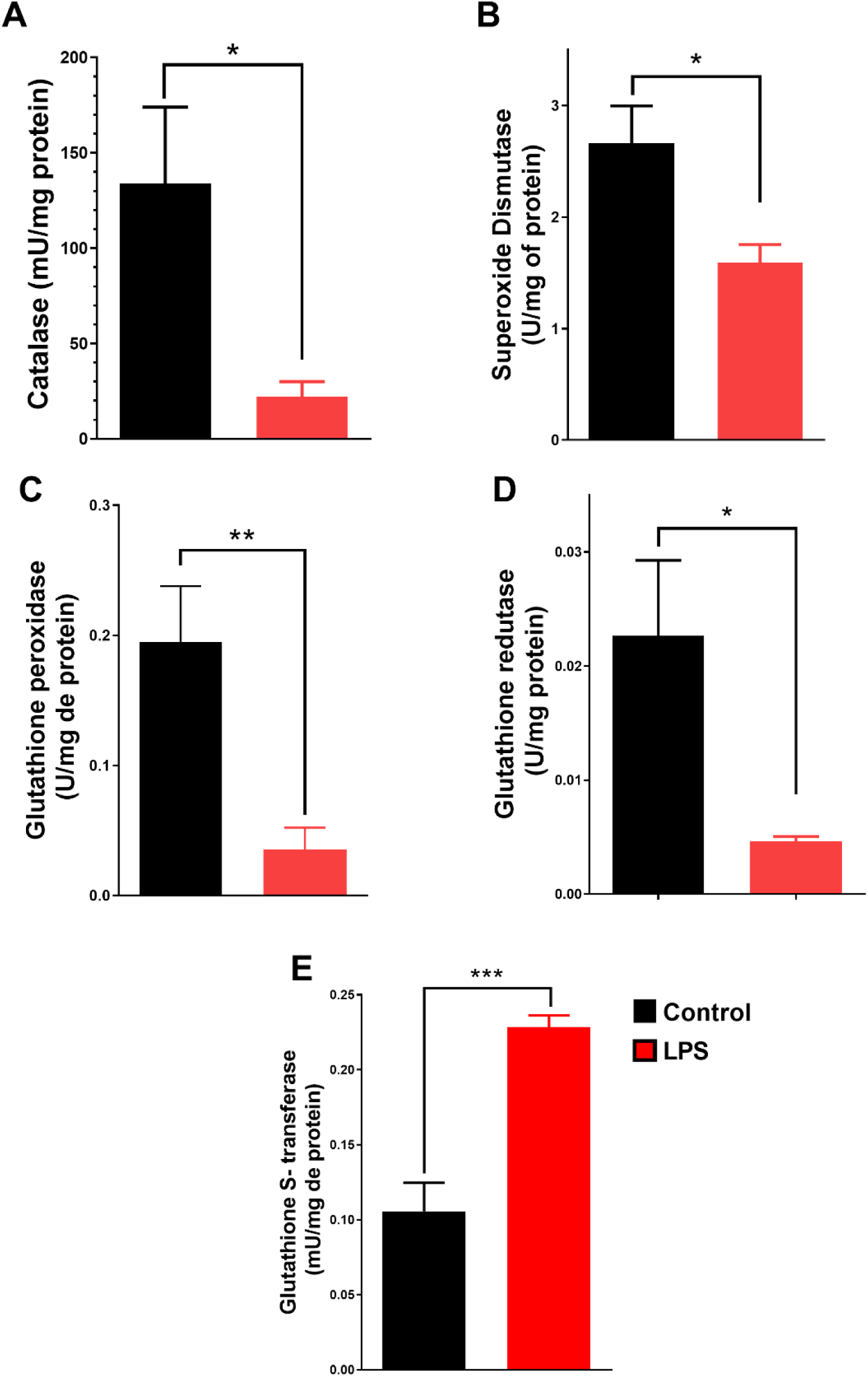
Cardiac antioxidant enzymes activity in LPS-treated rats. A, catalase activity. B, superoxide dismutase activity. C, glutathione peroxidase activity. D, glutathione reductase activity. E, glutathione S-transferase activity. (n = 4-5). * P < 0.05, ** P < 0.01, *** P < 0.001.

**Supplementary Fig. 4.**
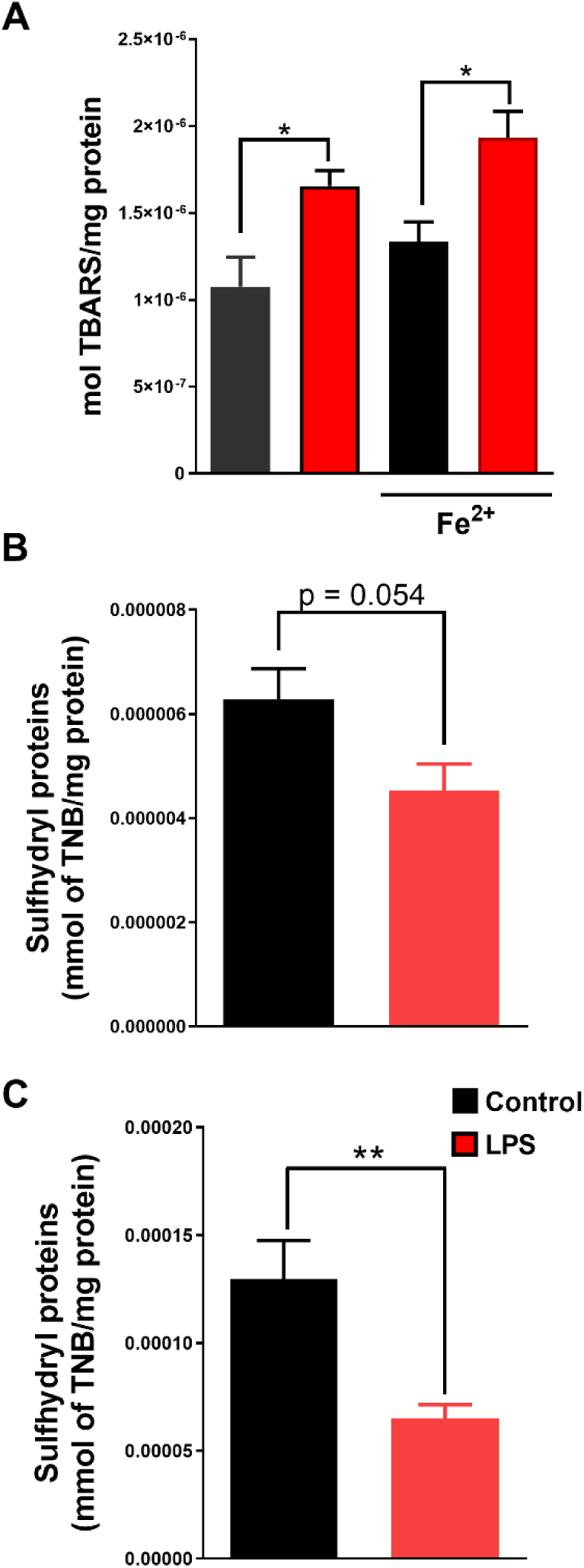
Effects of LPS-induced sepsis in lipid peroxidation and protein thiol oxidation in rat cardiac tissue. A, Lipid peroxidation (TBARS) increases in LPS-induced sepsis in the presence or absence of Fe^2+^. Mitochondrial (B) and cytosolic (C) protein sulfhydryl levels. (n = 4-5). * P < 0.05, ** P < 0.01.

